# Orthrus: an AI-powered, cloud-ready, and open-source hybrid approach for metaproteomics

**DOI:** 10.1101/2024.11.15.623814

**Authors:** Yun Chiang, Matthew James Collins

## Abstract

While metaproteomics provides invaluable insight into microbial communities and functions, significant bioinformatics challenges persist due to data complexity and the limitations of database searching. We introduce `Orthrus`, a hybrid approach combining transformer-based *de novo* sequencing (`Casanovo`) and database searching with rescoring (`Sage`+`Mokapot`). Benchmarking against PEAKS^®^11, `MaxQuant`, and `MetaNovo`, `Orthrus` demonstrates high peptide outputs, taxonomic diversity, and proteome coverage. `Orthrus` is Python-based and accessible to all via Google Colaboratory.

## Main

Microbial research has been propelled by large-scale initiatives, including multiple phases of Human Microbiome Projects (HMP ^1^ and iHMP ^2^) in the last two decades. Since coined in 2004 ^3^, metaproteomics has provided critical insight into the abundance, functions and composition of microbial communities. Despite the contributions, it appears that metaproteomics papers constitute only one percent of total proteomics submissions ^4^. Significant bioinformatics challenges, including the analysis of fragment ion spectra (MS2), have been cited as a major bottleneck ^5,6^. Microbiota are intrinsically diverse and vary in abundance; this poses challenges for selecting databases for peptide-spectrum matching. It is also evident that the use of a comprehensive but substantial database results in miscalculated false discovery rates (FDRs) and decreased peptide sensitivity ^7^. Indeed the NCBI nr has grown to 834 million entries for BLAST ^8^; this exemplifies the need to balance database comprehensiveness, computational efficiency and statistical quality controls in metaproteomics.

To mitigate the limitations of conventional database searching, hybrid, multi-step methods have been proposed to navigate database issues and improve protein identification. PEAKS^®^ is the state-of-the-art hybrid search engine that integrates *de novo* sequencing into conventional database searching. A recent iteration of PEAKS^®^ 11 leverages dynamic programming to shortlist proteins for database searching ^9^ and deep learning to boost identification. While PEAKS^®^ is a proprietary programme, `MetaNovo` offers an open-source approach combining sequence tagging, probabilistic ranking and `MaxQuant` ^10^. These hybrid strategies leverage *de novo* sequencing to improve the accuracy and sensitivity of database searching. However, hybrid approaches are often inaccessible due to associated licence fees and computational requirements. Accessibility and transparency are crucial for FAIR principles ^11^ and enabling high-throughput metaproteomics research.

Here we introduce `Orthrus`, a two-pronged system that makes a hybrid search strategy accessible via Google Colaboratory (Colab). `Orthrus` integrates `Casanovo`, a state-of-the-art transformer model that outperforms other neural networks in terms of amino acid and peptide precision ^12^, with `Sage` ^13^, a Rust-engineered search engine that prioritises processing speed using a fragment ion index popularised by `MSFragger` ^14^. `Orthrus` is AI-powered, cloud-ready and open-source that can be executed online in Colab or locally in a Python environment. It supports data dependent acquisition (DDA) experiments and standardised vendor-free instrument files. `Orthrus` is flexible and modular, adhering to the FAIR principles.

`Orthrus` (Fig. 1) loads vendor-free .mzML or .mgf files and calls `Casanovo` for *de novo* sequencing. Users can specify a model (.ckpt) and configuration file (.yaml) or use Casanovo’s default options. `Casanovo` outputs are filtered and concatenated together, dividing into overlapping tags. This mitigates the dominance of y-ions from the c-terminal ^15^, which may result in incorrect amino acid arrangements. The sizes of *de novo*-based tags are data-dependent, according to the median lengths of `Casanovo` predicted outcomes (len= 11 or 12, Supplementary Table 1). These *de novo*-based tags are subsequently string-matched with a large reference database for initial shortlisting. Matches are then evaluated and ranked using a Naïve Bayes Classifier. As a measure of conditional probability, the likelihood of a match depends on a spectral abundance factor and an intensity proxy. The spectral abundance factor represents the number of assigned spectra per protein, normalised by protein length ^16^, while the intensity proxy is based on the number of tryptic peptides generated *in silico* for each protein entry. The selection of these conditions are consistent with previous quantification workflows such as `MetaNovo` ^10^ and `IonQuant` ^17^. A confusion matrix and Receiver-operating characteristic curves (ROC) with cross-validation to evaluate the ranking model are provided in Supplementary Information. The most probable matches (≥ 95% probability) are shortlisted, and associated protein entries are returned as a single .fasta for database searching by `Sage`. `Mokapot` ^18^ rescores `Sage` .pin files using semi-supervised support vector machine (SVM) modelling implemented by `Percolator` ^19,20^. `Mokapot` is lightweight and flexible, supporting both static and joint rescoring models. Overall, `Orthrus` utilises a wide range of open-source Python tools, including but not limited to `Casanovo`, `Pyteomics`, `Biopython`, `Sage` and `Mokapot`. All these tools are preconfigured in two .ipynb notebooks and ready for online execution by Colab.

**Fig. 1.**
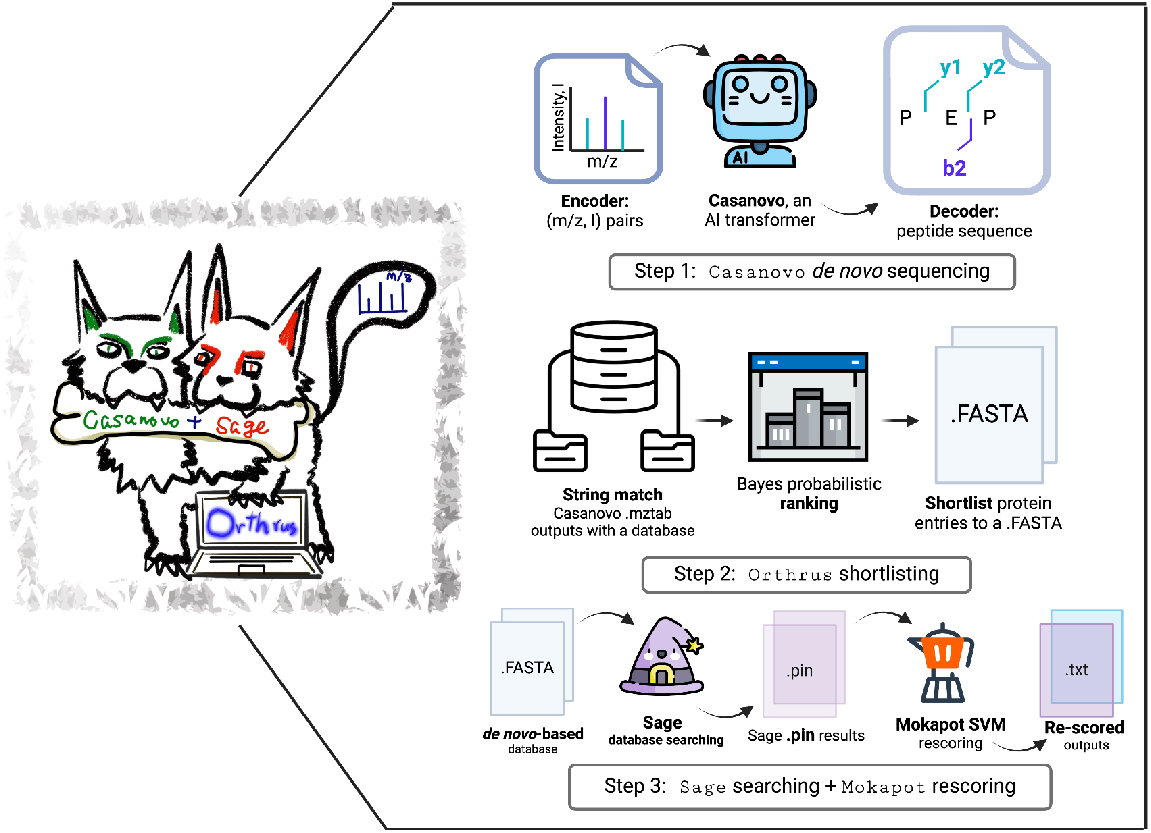
The architecture of `Orthrus`. It contains three key steps: (1) *de novo* sequencing powered by Casanovo’s transformer, (2) `Orthrus` shortlisting using Naïve Bayes classification, and (3) a *de novo*-first database for `Sage` searching and `Mokapo`t machine learning-based (support vector machine SVM) rescoring.

We benchmarked `Orthrus` against three search engines, including `MaxQuant`, `MetaNovo` and PEAKS^®^ 11. While `MaxQuant` is not a hybrid approach ^21^, it is included for its wide applications and is used as baseline data ^22^. The benchmarking dataset is circa one million mixed, high-impact modern human (PXD038906 ^23^) and ancient (PXD027613 ^24^) human coprolite spectra. The study of ancient microbes sheds light on microbial variations and long-term dynamics ^25^. The inclusion of ancient metaproteomics spectra is useful to test hybrid systems and how they deal with a wide range of low-abundance microbial proteins.

We compared the four search engines at three levels: peptide-spectrum matches (PSMs) with a FDR threshold of 0.01, the comparison of assigned peptides, taxonomic resolution, and host protein coverage (Fig. 2). At FDR 0.01, `Orthrus` (n=37,337) and PEAKS^®^ 11 (n=32,694) assigned most PSMs, which triple the outputs of `MetaNovo` (n=12,646) and quadrupole those of `MaxQuant` (n=8,545). The UpSet plot illustrates a considerable agreement between the four search engines. `Orthrus` and PEAKS^®^ 11 again identified most peptides. Given the similar performance between `Orthrus` and PEAKS^®^ 11, we further interrogated the taxonomic assignments using `UniPept` ^26^. It appears that a wide range of microbial phyla was identified by `Orthrus`, while PEAKS^®^ 11 generated more non-specific (root) peptides. Lastly, `Orthrus` achieves the highest modern (18.60%) and ancient (15.11%) host protein sequence coverage. This measurement is the mean sequence coverage of all identified ancient and modern human proteins by each search engine. A non-parametric Kruskal-Wallis test and post-hoc Dunn’s comparisons also support that these `Orthrus` proteome coverage results are statistically significant from the other 3 engines (Supplementary Table 2).

**Fig. 2.**
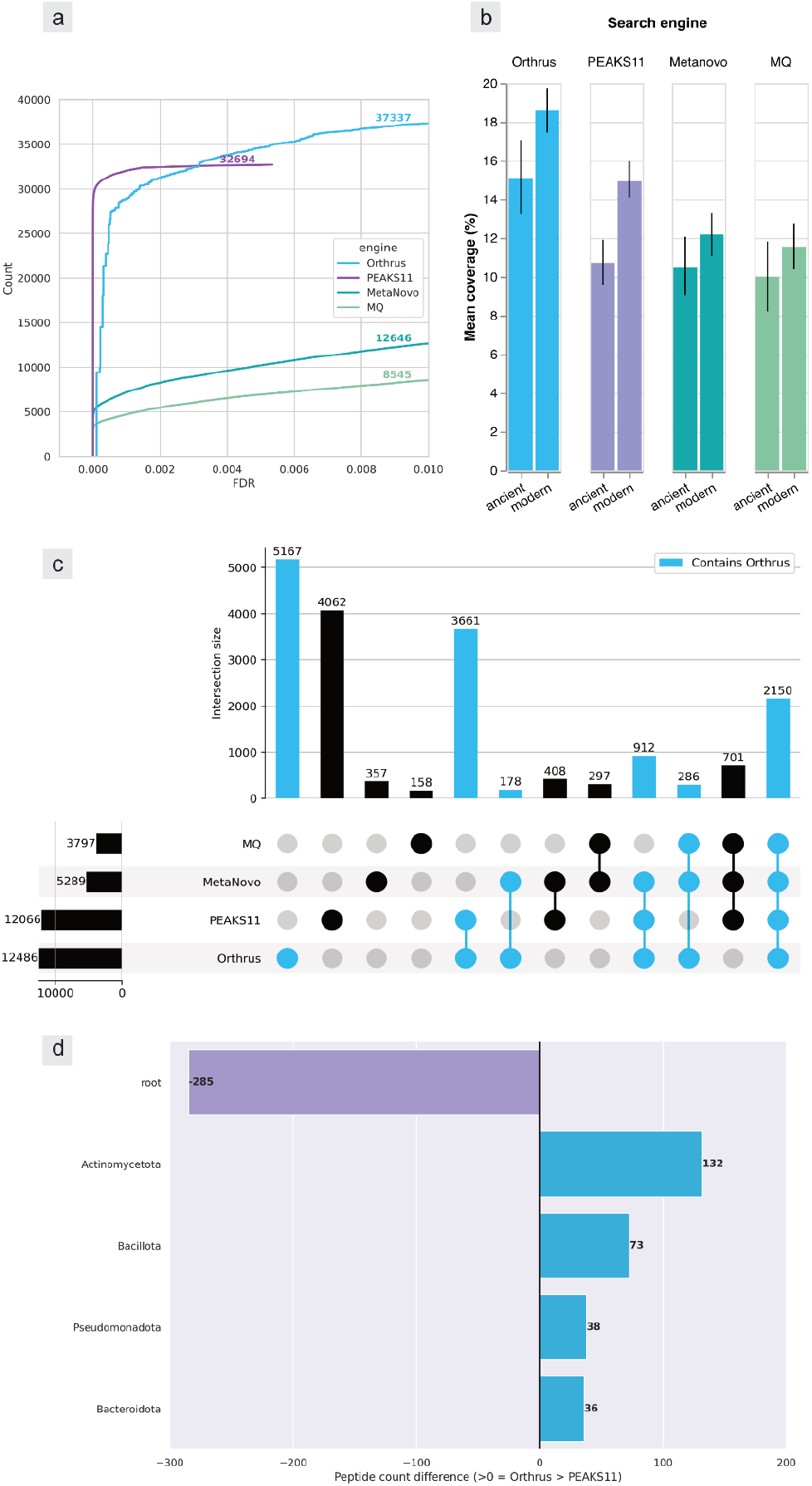
(a) The trends and numbers of PSMs below FDR 0.01 for the four search engines. (b) Bar plots with error bars to illustrate the mean coverage (%) of the human host proteome identified by each engine. (c) An UpSet plot highlighting the intersection of identified peptides, with `Orthrus` results coloured in light blue. (d) A diverging bar illustrating the combined (modern + ancient) peptide count differences between `Orthrus` and PEAKS^®^ 11 at the phylum level. These microbial phyla were all reported in the original modern metaproteomics publication^23^.

To demonstrate the scalability of `Orthrus`, we analysed all modern coprolite data (= 9,047,508 spectra) from PXD038906, and the entire reviewed Swiss-Prot and automated TrEMBLE databases evident at protein/transcription levels (n=2,207,956) were used for shortlisting. At the threshold of FDR 0.01, `Orthrus` identified 486,211 PSMs and 360,494 peptides, assigned to 1665 lowest common ancestors (LCAs) by `UniPept`. We used the host abundance (genus=*Homo*) as the baseline and illustrated the most abundant taxa (log_2_ fold change ≥ -4) at the genus level in the human coprolite samples. A wide range of dietary (plant + animal, 62.96%) and microbial (associated with the gut or faeces, 37.04%) proteins are present (Supplementary Fig. 3). We further analysed the functions of these abundant taxa using GO annotations. It is evident that host proteins contribute to immune responses, and dietary proteins are associated with seed metabolism and proteolysis. Microbes are also involved in a variety of translational and metabolic processes. Only human proteins were searched and microbes were analysed at the phylum level for multi-omics in the original publication. We demonstrated that `Orthrus` expanded taxonomic assignment and functional analysis beyond host proteins with confidence and high resolution.

While AI-powered de novo sequencing lacks innate FDR measurements, we propose a two-pronged `Orthrus` approach which leverages a transformer model while maintaining controls for challenging metaproteomics datasets. `Orthrus` is accessible and scalable, outperforming established search engines in taxonomic diversities and protein coverage.

## Methods

### Running Orthrus

`Orthrus` is available as preconfigured Jupyter notebooks suitable for any notebook environments, for example, JupyterLab, Jupyter Notebook via Anaconda, and Google Colab. Orthrus is optimised for Colab and ready for execution online, with tutorials available on Github (https://github.com/yc386/orthrus_metaproteomics).

### Colab notebooks

`Orthrus` has three annotated notebooks for users. The first notebook utilises Colab GPU runtime to accelerate *de novo* sequencing via a built-in Nvidia^®^ CUDA^®^ driver (version 12.2, continually updated by Colab). An A100 GPU runtime (GPU RAM: 40 GB, system RAM: 83.5 GB) is recommended for optimal performance, but a T4 runtime (GPU RAM: 15 GB, system RAM: 51 GB) is accessible to all Colab users. The second notebook leverages Colab TPU runtime, ideal for Sage and database searching because of the CPU-intensive nature and a potential high RAM requirement. A TPU runtime (40 CPU cores, 334.6 GB RAM) is available without a Colab subscription. The third notebook provides an example of using customised `Scikit-learn` inputs for `Mokapot` rescoring.

### Orthrus implementation

`Orthrus` integrates a wide range of open-source, Python-based tools: `Pandas`, `Numpy`, `Joblib`, `Pyteomics`, `Biopython`, `Scikit-learn`, `Casanovo`, `Sage`, and `Mokapot`. All libraries are installed by `PyPI` and latest versions are fetched by default. `Sage` is implemented through `Bioconda`. `Orthrus` consists of three key steps: *de novo* sequencing (`Casanovo`), Orthrus shortlisting, and database searching (`Sage`) with rescoring (`Mokapot`).

### *de novo* sequencing

`Casanovo` is executed in Colab through a command line interface. `Orthrus` supports user-defined configuration files (.yaml) and models (.ckpt). If unspecified, `Casanovo` default settings are used. `Casanovo` provides both tryptic and non-enzymatic models (fine-funed with different enzyme inputs) without retraining ^27^.

### Orthrus shortlisting

`Casanovo` outputs (.mztab) are parsed and filtered by the maximum value of search_engine_score [1] (as part of `Casanovo` prediction outputs) below zero. As outlined in the publication ^12^, a prediction is marked negatively if the predicted mass of a sequence deviates beyond its observed precursor mass and the defined mass tolerance range.

Filtered `Casanovo` predictions without PTMs are concatenated and divided into overlapping sequence tags, which are string matched with a user-provided reference database. We have also included a default option to fetch the latest Swiss-Prot database (uniprot_sprot.fasta) from the UniProt FTP server. Since `Casanovo` does not differentiate between leucine (L) and isoleucine (I), L is equated with I in the provided database.

Database sequences are divided into overlapping tags in the same manner as *de novo* tags. The sizes of both *de no* and database tags are based on the median lengths of `Casanovo` predictions (Supplementary Table 1). For *de no* and database tags to be matched, all amino acid sequences must be identical. A matched protein entry class is assigned if it contains at least two matched *de novo* tags.

After string matching, Bayes’ theorem is used to shortlist protein matches based on conditional probabilities:

P (A|B)=[ P (B|A)* P(A)] / P(B), where an event A is a matched protein entry (containing ≥ 2 matched *de novo* tags), and B events represent conditions, including a spectral abundance factor (SAF) and an intensity proxy.

The SAF is calculated for each protein entry based on the number of *de novo* matches normalised by protein length, consistent with MetaNovo’s implementation ^10,16^. Moreover, the intensity proxy is approximated by a ratio of tryptic peptides to the total number of sequence tags per database protein entry. Tryptic peptides are simulated in *silico*, cleaving at arginine and lysine without proline restriction. The two conditions contain continuous values and are independent of each other. They are normalised (MinMaxScaler`, Scikit-learn)` before Naive Bayes classification. Most probable matches (≥ 95% probability) are shortlisted and exported as a .fasta file for input into `Sage`.

### Sage database searching + Mokapot rescoring

`Orthrus` automatically updates `Sage` configuration (.json) based on user inputs. If unspecified, `Mokapot` is executed automatically after `Sage` to rescore its search output (.pin). We have included different model options and parameter settings for `Mokapot`: `Percolator` -based SVM and non-linear gradient boost classifier based on a published `XGBoost` implementation ^18^. A separate Jupyter notebook is available as an example of utilising a user-defined `Scikit-learn` model for `Mokapot`. Rescoring can be done individually for each experiment, or a joint model could be learned from all matches across experiments, according to `Mokapot`’s implementation ^18^.

### Benchmarking

To benchmark Orthrus, modern human metaproteomics data were downloaded from PXD038906 ^23^, of which 591,035 spectra were randomly selected. Additionally, ancient human coprolite data was collected from PXD027613 ^24^, and all 561,120 ancient MS2 spectra were included. In total the benchmarking dataset consisted of 1,152,155, mixed, modern and ancient MS2 spectra for benchmarking. The collected .RAW files were converted to .mzML or .mgf (for MetaNovo only) files by `ThermoRawFileParser` (1.7.3) ^28^ using default settings without file compression.

The ∼1.15 mixed MS2 spectra were searched using PEAKS^®^ 11 (11.5 build 20231206), `MaxQuant` (2.5.2.0), `MetaNovo` (Galaxy Version 1.9.4+galaxy4), and `Orthrus` (1.0.0). `ThermoRawFileParser`, PEAKS^®^ 11, and `MaxQuant` were executed on a local workstation equipped with 18 CPU cores (Intel^®^ Xeon^®^ W-2295) with 256 GB RAM. `MetaNovo` was carried out by the European galaxy server and `Orthrus` was run on Google Colab Pro+ (A100 GPU runtime, 83.48 GB RAM).

The protein sequence database used in the benchmarking test consisted of the reviewed human and bacterial proteins from Swiss-Prot (2024-03 release), and common contaminants (total n=356,839 entries without duplicates). For all searches, tryptic digestion without proline suppression was selected. Two missed cleavages were allowed. The carbamidomethylation of cysteine was set as a fixed modification. For PXD038906 (modern), the variable modifications were N-terminal acetylation and the oxidation of methionine. As for PXD027613 (ancient), deamidation (asparagine + glutamine) and the oxidation of methionine were used. Up to five modifications were allowed per identification The precursor mass tolerance was 7 ppm (consistent with `MaxQuant` optimised settings for Orbitrap™ instruments), and the fragment mass tolerance was 0.02 Da.

Given the nature of ancient metaproteomics samples, adjustments were made for `Orthrus` (1.0.0). The non-enzymatic model was used for all the ancient MS2 spectra during `Casanovo` *de novo* sequencing, since they may contain unspecific cleavage due to protein degradation. A joint model was applied to rescore the ancient search results using `Mokapot`, due to low PSM outputs per experiment. Joint modelling is able to mitigate the limitations of using static modelling for low-abundance single cell proteomics datasets with few PSMs ^18^. In contrast, a tryptic model was used for *de novo* searching the modern MS2 spectra. A separate model was used to rescore each modern metaproteomic experiment, since it was not constrained by low PSM outputs. All other parameters were consistent with other search engines.

Due to the complexity of metaproteomics datasets and the presence of conserved sequences (shared across different species), `UniPept` (6.0, web version) ^29^ and its lowest common ancestor algorithm (LCA) were used for taxonomic assignments. This latest version of `UniPept` also supports the metaproteomics analysis of peptides with missed cleavages, which is ideal for ancient and semi-tryptic peptides caused by protein degradation over time.

### Evaluating Naïve Bayes classification

The effectiveness of Naïve Bayes classification was evaluated by accuracy, precision, F1 scoring, a confusion matrix and ROC curves with five-fold cross-validation. All evaluation metrics were obtained from `Scikit-learn` (1.5.2). Detailed plots are reported in Supplementary Figs 1& 2. The data used for the evaluation was the ∼ one million benchmarking dataset. We employed the five-fold cross-validation approach by splitting the benchmarking data into five equal parts, using a different (20%) fraction for testing and the remaining (80%) for training during each of the five iterations.

Overall, the Bayes classification model demonstrated high accuracy (99.02 %), precision (82.09 %), recall (99.48 %) and f1 scoring (89.95 %). The confusion matrix shows that true negatives (98.99 %) and positives (99.48 %) dominate each class. The percentage was calculated per class, for example, for class ==1, the ratio was the number of true positives to total predicted positives. The ROC curve shows that on average, the area under curve (AUC) is 0.9948, representing a 99.48% chance of correctly labelling a random input.

### Scalability test

A large and comprehensive reference database is often required to accommodate the complexity of metaproteomics samples. Scalability is essential for the development of metaproteomics bioinformatics pipelines.

All 9,047,508 modern human stool MS2 spectra from PXD038906 were included, and reviewed Swiss-Prot and automated TrEMBLE at protein/transcription levels (total entries=2,207,956, 2024-04 release) were used for `Orthrus` (1.0.0). The tryptic model was used for `Casanovo` *de novo* searching. For `Sage` searches, parameters were consistent with the benchmarking with slight adjustments to accommodate the larger database. Only three modifications were allowed per assignment. A separate model was used to rescore `Sage` search per experiment using `Mokapot`. Taxonomic assignments were also carried out by `UniPept` (6.0, web version).

### Ancient protein authentication

Since ancient proteins are susceptible to contamination, authentication is required to avoid misinterpretations and ensure research rigour. We used `Anubis` (1.0.0) (https://github.com/yc386/anubis_palaeoproteomics), an open-source Python tool for authenticating ancient proteins, to calculate deamidation rates for the ancient MS2 dataset and compared the patterns of putative ancient proteins with trypsin processed in the same batch (Supplementary Fig. 4).

## Data & Code availability

Tutorials and pre-configured, ready-for-execution Jupyter Notebooks are available on Github (https://github.com/yc386/orthrus_metaproteomics). The Github repository also contains all data and code to reproduce statistical analysis and figures in the study.

## Supplementary Information

**Supplementary Table 1:**
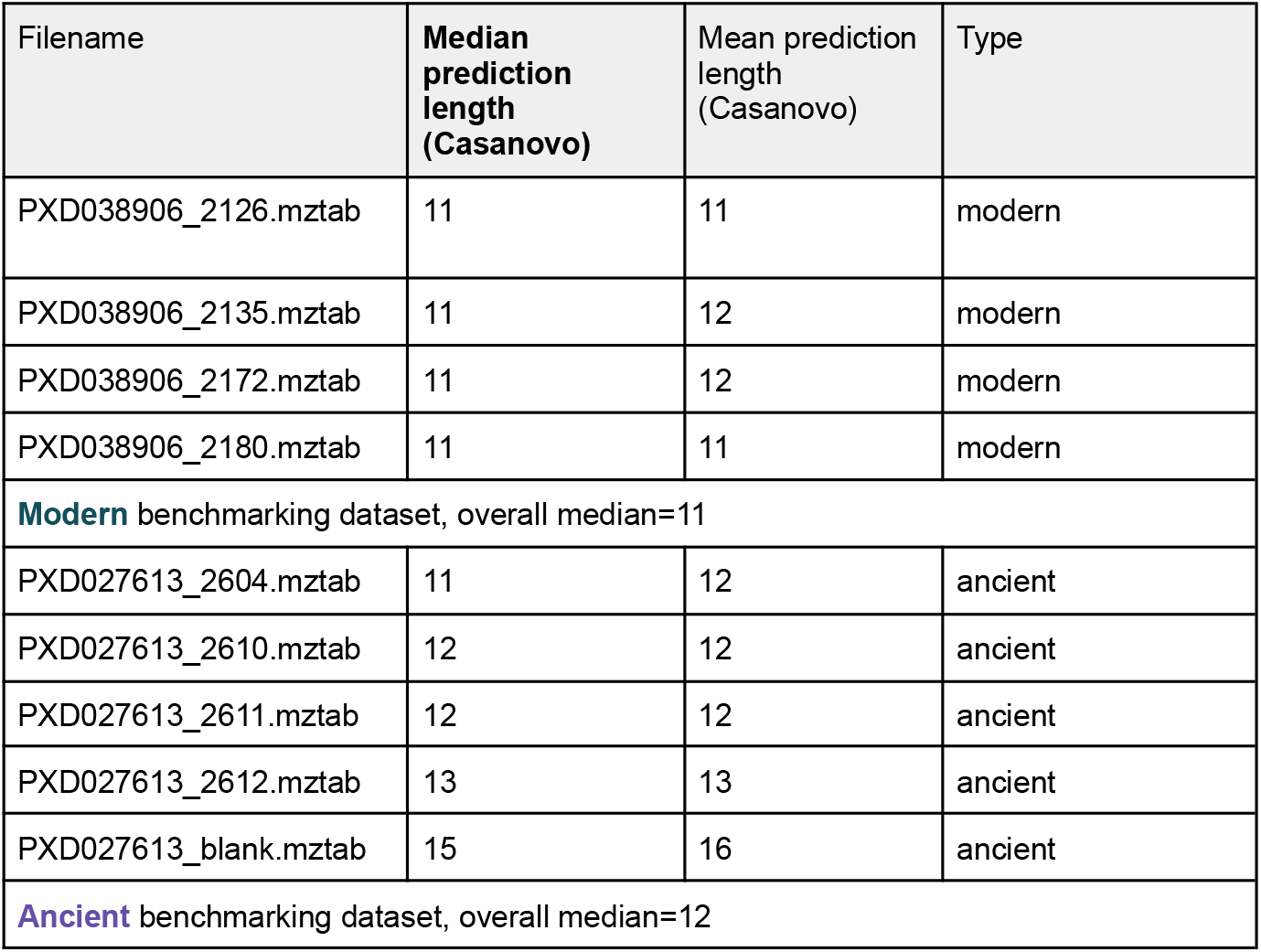
The median and mean lengths of Casanovo sequence predictions were calculated for each output file. The overall median lengths were used to determine the size of de novo sequence tags for modern (len=11) and ancient (len=12) benchmarking datasets.

| Filename | Median prediction length (Casanovo) | Mean prediction length (Casanovo) | Type |
| --- | --- | --- | --- |
| PXD038906_2126.mztab | 11 | 11 | modern |
| PXD038906_2135.mztab | 11 | 12 | modern |
| PXD038906_2172.mztab | 11 | 12 | modern |
| PXD038906_2180.mztab | 11 | 11 | modern |
| **Modern** benchmarking dataset, overall median=11 |  |  |  |
| PXD027613_2604.mztab | 11 | 12 | ancient |
| PXD027613_2610.mztab | 12 | 12 | ancient |
| PXD027613_2611.mztab | 12 | 12 | ancient |
| PXD027613_2612.mztab | 13 | 13 | ancient |
| PXD027613_blank.mztab | 15 | 16 | ancient |
| **Ancient** benchmarking dataset, overall median=12 |  |  |  |

**Supplementary Fig. 1:**
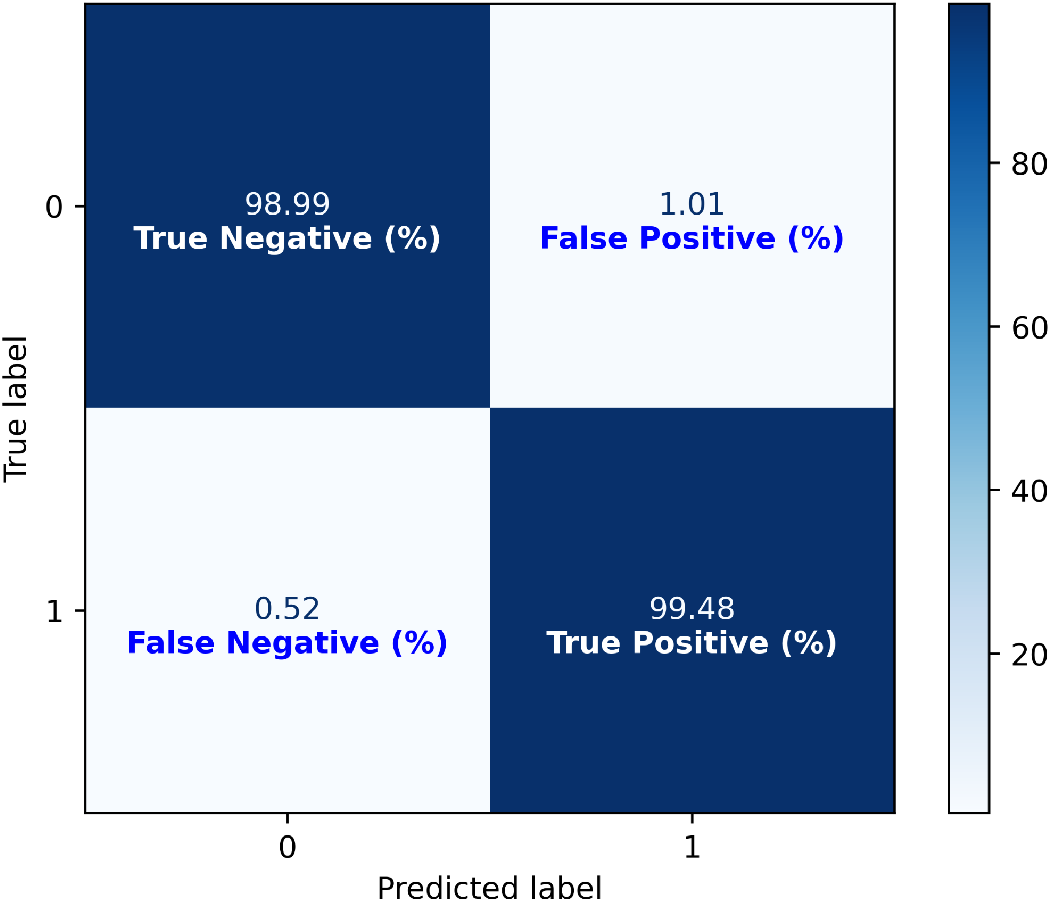
A confusion matrix for evaluating the Naïve Bayes classification model. The matrix visualises true negatives, false positives (Type I errors), false negatives (Type II errors), and true positives. The c. 1 million, mixed benchmarking dataset was used to build the confusion matrix. Percentages were calculated per class (0: unmatched, 1:matched). Overall, both false positives and negatives were low.

**Supplementary Fig. 2:**
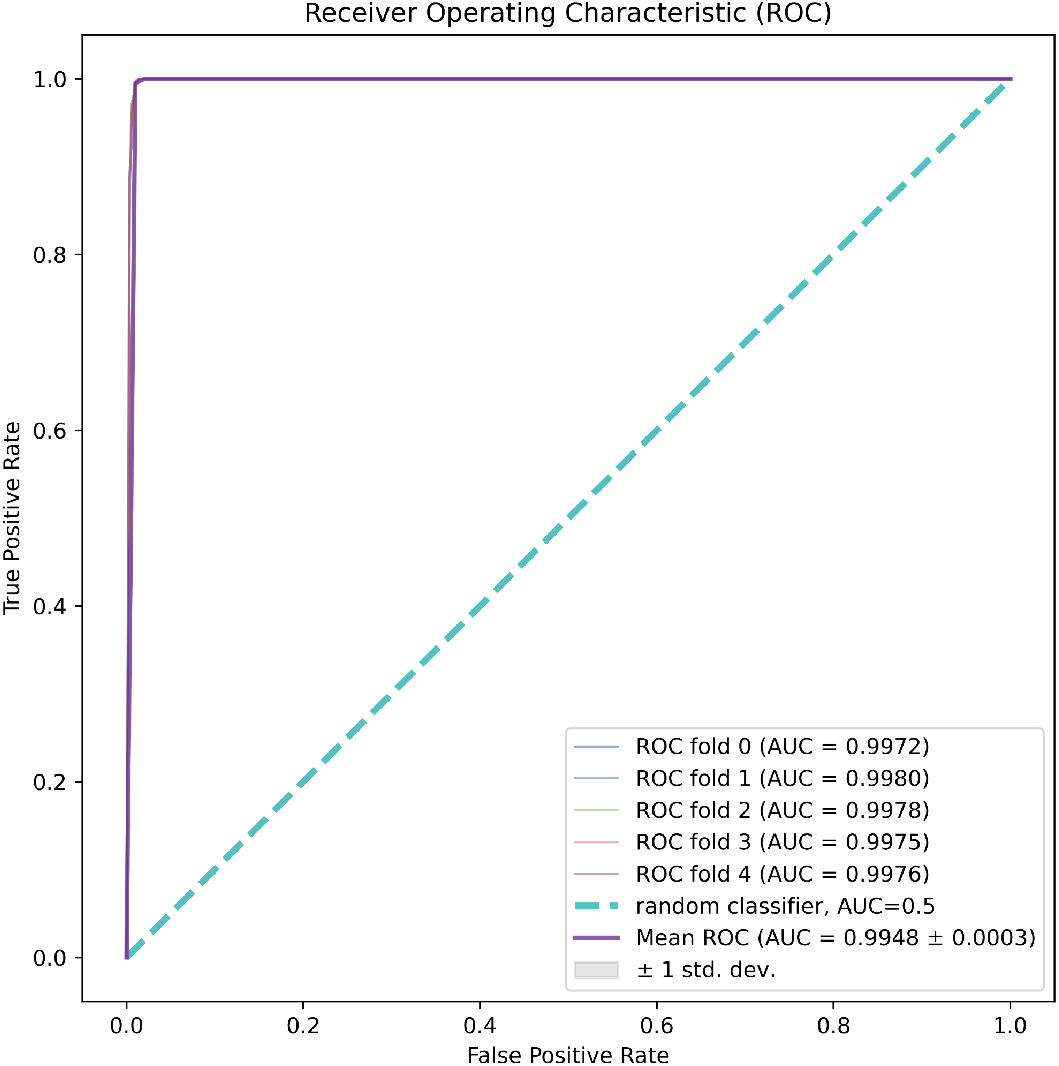
A ROC curve for evaluating the Naïve Bayes binary classification with five folds of cross evaluation. True positive rate was plotted against false negative rate. A random classifier (AUC=0.5) was highlighted. It is clear that the AUC of each fold and the mean are all higher than the random classifier. The benchmarking dataset was used to conduct the ROC analysis.

**Supplementary Table 2:** Sequence coverage was calculated for human protein entries as part of the benchmarking outputs to estimate host proteome coverage. Common contaminants were excluded. A non-parametric Kruskal-Wallis was conducted for all four search engines using Scipy. Kruskal-Wallis H statistics and *p* values were reported in the table for modern and ancient human host proteomes. As these *p* values are significant, post hoc Dunn’s tests were performed (by scikit-posthocs) to evaluate pairwise significance with a *p* value adjusted by the Bonferroni correction. It is evident that Orthrus results are significantly different from the other search engines for both modern and ancient human proteomes.

| Modern host proteome |  |  |  |  |
| --- | --- | --- | --- | --- |
| Kruskal-Wallis H statistic: 129.3491704393071, $p$ value=7.470090196236122E-28 | | | | |
| Post hoc Dunn's tests with Bonferroni correction |  |  |  |  |
|  | Orthrus | PEAKS11 | MetaNovo | MaxQuant |
| Orthrus | 1.000000 | 1.942014E-08 | 1.926547E-20 | 3.068539E-21 |
| PEAKS11 | 1.942014E-08 | 1.000000 | 7.333023E-05 | 7.435320E-06 |
| MetaNovo | 1.926547E-20 | 7.333023E-05 | 1.000000 | 1.000000 |
| MaxQuant | 3.068539E-21 | 7.435320E-06 | 1.000000 | 1.000000 |

**Supplementary Table 2:** Sequence coverage was calculated for human protein entries as part of the benchmarking outputs to estimate host proteome coverage.
| Ancient host proteome |  |  |  |  |
| --- | --- | --- | --- | --- |
| Kruskal-Wallis H statistic: 27.277599998376033, $p$ value=5.1488875361284335E-06 | | | | |
| Post hoc Dunn's tests with Bonferroni correction |  |  |  |  |
|  | Orthrus | PEAKS11 | MetaNovo | MaxQuant |
| Orthrus | 1.000000 | 0.000017 | 0.000199 | 0.000339 |
| PEAKS11 | 0.000017 | 1.000000 | 1.000000 | 1.000000 |
| MetaNovo | 0.000199 | 1.000000 | 1.000000 | 1.000000 |
| MaxQuant | 0.000339 | 1.000000 | 1.000000 | 1.000000 |

**Supplementary Fig. 3.**
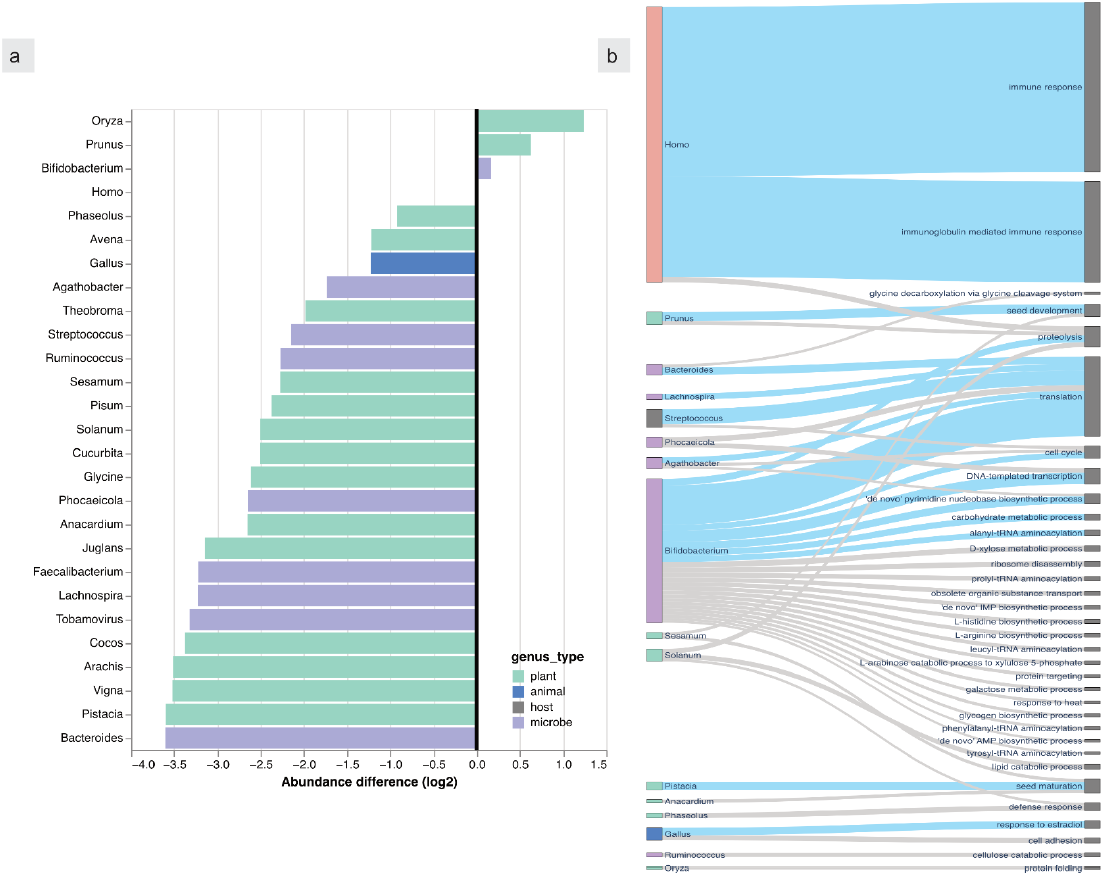
(a) A diverging bar plot highlighting the most abundant taxa at the genus level. The abundance (based on MS2 fragment ion intensities) of the host (human, *Homo sapiens*) was used as the baseline. Positive differences (log_2_ values) indicate higher abundance than the host. Horizontal bars were coloured by four broad taxonomic groups (plant/animal/host/microbe). (b) A Sankey diagram showing the relationship between abundant taxa at the genus level and most common functions (top 50).

**Supplementary Fig. 4.**
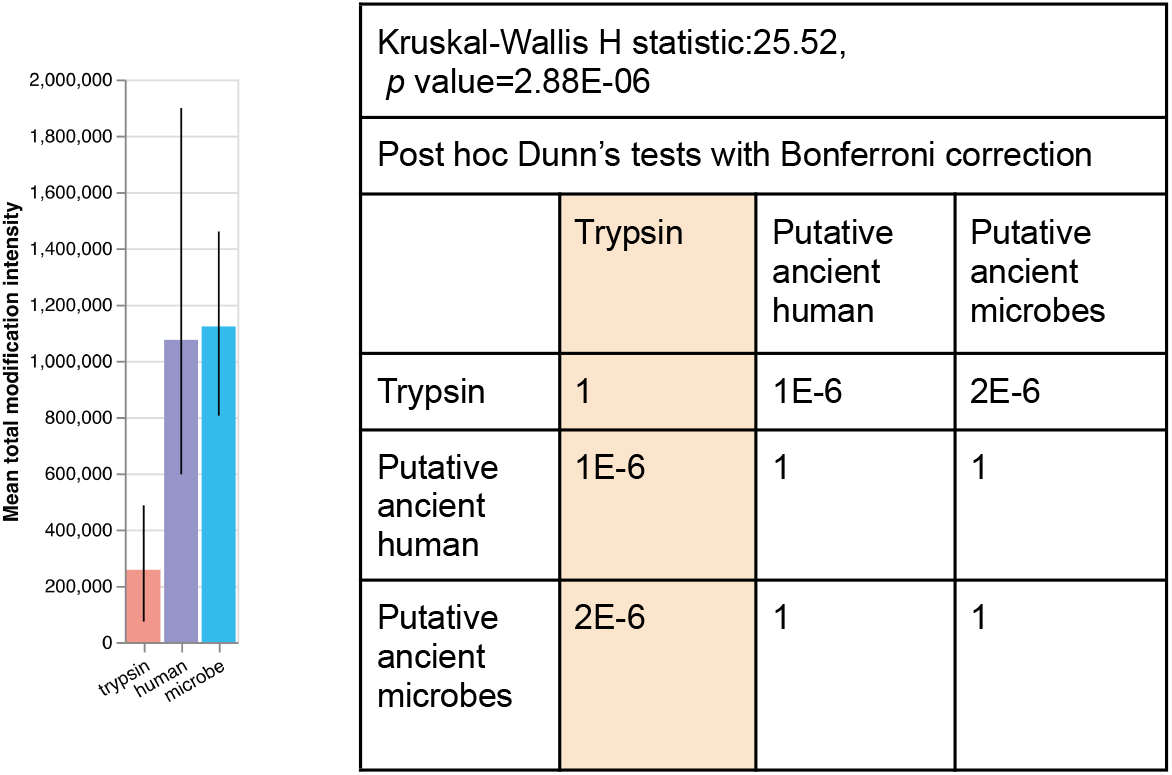
The mean and total modification intensity calculated by `Anubis`. Briefly, the MS2 intensities of oxidation (methionine) and deamidation (asparagine + glutamine) were estimated at the amino acid position level and extrapolated to the protein level. Glutamine deamidation intensities were weighted to reflect its time-dependent nature. Human-related common contaminants (human keratins) were excluded from the putative ancient human protein group. The mean modification intensities of putative ancient human and microbial proteins are significantly different from the modification pattern of trypsin processed with the ancient samples, corroborated by a non-parametric Kruskal-Wallis test and post hoc Dunn’s comparisons with the Bonferroni correction.

